# Comparison of Mucosal and Intramuscular Immunization against SARS-CoV-2 with Replication-Defective and Replicating Single-cycle Adenovirus Vaccines

**DOI:** 10.1101/2021.04.20.440651

**Authors:** Haley E. Mudrick, Erin B. McGlinch, Brian J. Parrett, Jack R. Hemsath, Mary E. Barry, Jeffrey D. Rubin, Chisom Uzendu, Michael J. Hansen, Courtney L. Erskine, Virginia P. VanKeulen, Aleksandra Drelich, Chien-Te Kent Tseng, Shane Massey, Madiha Fida, Gina A. Suh, Tobias Peikert, Matthew S. Block, Gloria R. Olivier, Michael A. Barry

## Abstract

SARS-CoV-2 enters the body at mucosal surfaces, such as the nose and lungs. These events involve a small number of virions at these mucosal barriers and are therefore a strategic point to stop a COVID-19 infection before it starts. Despite this, most vaccines against COVID-19 are being injected into the muscle where they will not generate the highest levels of mucosal protection. The vaccines that are approved for use in humans are all replication-defective (RD) mRNA, DNA, or adenovirus (Ad) vaccines that do not amplify antigen transgenes. We developed single cycle adenovirus (SC-Ad) vectors that replicate antigen genes up to 10,000-fold in human cells, but that are disabled from producing infectious Ad particles. We show here that SC-Ad expressing the full-length SARS-CoV-2 spike protein produces 100-fold more spike protein than a matched RD-Ad-Spike vector. When Ad-permissive hamsters were immunized with these vaccines by intranasal (IN) or intramuscular (IM) routes, SC-Ad produced significantly stronger antibody responses as compared to RD-Ad against the spike protein that rose over 14 weeks after one immunization. Single IN or IM immunizations generated significant antibody responses in serum and in bronchoalveolar lavages (BALs). IN priming, but not IM priming, generated HLA-restricted CD8 T cell responses in BALs. SC-Ad-Spike generated antibodies that retain binding to spike receptor binding domains (RBDs) with mutations from new viral variants. These data suggest empowering the genomes of gene-based vaccines with the ability to amplify antigen genes can increase potency. This may be particularly advantageous when applying mucosal vaccines to combat mucosal pathogens like SARS-CoV-2.

**One Sentence Summary:** Arming adenovirus vaccines with the ability to replicate vaccine antigen genes may increase potency for systemic, or more importantly, mucosal immunization against mucosal pathogens.

## Introduction

In December 2019, a cluster of pneumonia cases were identified in Wuhan, China, which were later found to be caused by a novel coronavirus: severe acute respiratory syndrome coronavirus-2 (SARS-CoV-2) (Li & al., 2020) (Gralinski & Menachery, 2020). As of April 2021, there have been over 130,000,000 cases and nearly 3,000,000 deaths world-wide (Johns Hopkins University Coronavirus Resource Center).

Nearly every vaccine technology has been deployed to combat this pandemic (reviewed in (*1*)). mRNA vaccines advanced through the development and regulatory processes most quickly and have been given emergency authorization from the FDA and other international regulators. Replication-defective adenovirus (RD-Ad) gene-based vaccines have also been advanced by several countries and companies, including chimpanzee Ad, human Ad serotype 5 (Ad5), human Ad serotype 26 (Ad26), and others. Each of these vaccines has their strengths and weaknesses, and most will not be revealed until human studies are completed.

While many of the advanced vaccines have great promise, they perhaps miss out on two opportunities to combat SARS-CoV-2 and other mucosal pathogens. First, most COVID-19 vaccines are administered intramuscularly (IM) and not at the site of SARS-CoV-2 entry into the body. Second, most COVID-19 vaccines do not harness the power of transgene replication to amplify antigen production and immune responses.

Most Ad vaccines are RD-Ads (reviewed in (*2, 3*)). For example, the Johnson & Johnson human Ad26 vaccine (*4*), the ChAdOx1 vaccine from Oxford/AstraZeneca (*5*), the Russian Sputnik V Ad26 and Ad5 vaccines (*6*), and most others are RD-Ad vaccines. Converting a wild replication-competent Ad to an RD-Ad is achieved by deleting the adenovirus’ pivotal master regulator gene, E1, to prevent them from causing Ad infections. An RD-Ad can efficiently deliver vaccine genes into cells to transcribe and translate vaccine antigens. However, after gene delivery, the DNA genome of an RD-Ad is not replicated (reviewed in (*2, 3*)). Therefore, one incoming RD-Ad vaccine antigen gene remains one gene. This gene can be very efficiently expressed, but it is not amplified.

By contrast, an E1-intact replication-competent adenovirus (RC-Ad) vector will infect a human cell and replicate an antigen gene DNA up to 10,000-fold in each infected cell (*7–18*). While RC-Ad vaccines are documented to be more potent than benchmark RD-Ad vectors, RC-Ads can cause actual adenovirus infections in vaccinees (*19*).

We developed single-cycle Ad (SC-Ad) vectors to take advantage of DNA replication while disabling the production of infectious progeny viruses (*20–23*). SC-Ads retain E1 genes and replicate their DNA just as well as RC-Ads, but are deleted for the gene for Ad’s pIIIa capsid cement protein, so that they produce empty defective particles (*20*). SC-Ads appear able to generate more robust and persistent immune responses than RD-Ads (*21, 24*) and have shown promise as vaccines against influenza (*24*), Ebola virus, HIV-1 (*25–27*), and against *Clostridium difficile* after single immunization (*22, 28*).

In this work, we generated RD-Ad and SC-Ad vectors expressing the wild-type original SARS-CoV-2 spike protein. Here, we compare the ability of RD-Ad and SC-Ad to produce the spike protein and generate immune responses in small animals. We also compare the ability of mucosal intranasal (IN) immunization relative to systemic intramuscular (IM) immunization to generate immune responses in systemic and mucosal compartments.

## Materials and Methods

### Single-cycle Adenovirus Expressing Wild-type SARS-CoV-2 spike

A codon-optimized cDNA encoding the original wild-type spike protein from severe acute respiratory syndrome coronavirus 2 isolate 2019-nCoV_HKU-SZ-002a_2020, accession number MN938384.1 was synthesized by Genewiz. This full-length sequence was inserted into the shuttle plasmid pAd6-NdePfl-CMV-MCS-3X-LZL. This sequence was recombined into pAd6-ΔE1-ΔE3 and pAd6-ΔIIIa-ΔE3 by red recombination as in (*20–22*) to generate RD-Ad6-spike and SC-Ad-Spike, respectively. These viruses were cut with AsiSI to liberate their viral genomes, and these were transfected into 293-IIIA cells to rescue the viruses. The viruses were purified from 10 Plate CellStacks (Corning) on two CsCl gradients and used as virus particles (vp) based on OD260 measurements (*20–22*).

### Western Blotting

Human A549 lung cells were infected with RD- or SC-Ad spike at the indicated multiplicities of infection (MOI) and harvested 24 hours later. 5X sodium dodecyl sulfate with 5mM dithiothreitol was added to cell lysate and heat-inactivated at 95°C for 5 minutes. Cell lysate was run by western blot using PowerPac™ HC (Bio-Rad) at 110 volts for 70 minutes. Gel was incubated in 1X transfer buffer while the membrane was prepared. Membrane was prepared by soaking 15 seconds in methanol, shaking 2 minutes in ddH2O on orbital shaker, and shaking 5 minutes in 1X transfer buffer. Gel was transferred to membrane via TransBlot® SD Semi-Dry Transfer Cell (Bio-Rad) at 15 volts for 15 minutes. Membrane was washed with 1X TBST on orbital shaker for 5 minutes. Membrane was incubated on orbital shaker in blocking buffer (5% milk powder in TBST) for 2 hours at room temperature. Membrane was washed with 1X TBST 3 times for 15 seconds each, then 3 times for 5 minutes each. Membrane was incubated in primary antibody, SARS-CoV-2 spike antibody [1A9] (GeneTex), diluted in blocking buffer at a 1:1000 dilution for 1 hour. Membrane was washed with 1X TBST 3 times for 15 seconds each, then 3 times for 5 minutes each. Membrane was incubated in secondary antibody, GOXMO HRP HIGH XADS (Invitrogen), diluted in blocking buffer at a 1:10,000 dilution for 1 hour. Membrane was washed with 1X TBST 3 times for 15 seconds each, then 3 times for 5 minutes each. Membrane was coated with 750 μL SuperSignal™ West Maximum Sensitivity Substrate (Thermo Scientific) and imaged on the ChemiDoc™ Imaging System (Bio-Rad).

### Animals

BALB/c mice were purchased from Charles River Laboratories. Syrian hamsters were purchased from Envigo. All animal handling and experiments were carried out according to the provisions of the Animal Welfare Act, PHS Animal Welfare Policy, the principles of the NIH Guide for the Care and Use of Laboratory Animals, and the policies and procedures of the Mayo Clinic. This study was conducted in Mayo Clinic’s AAALAC (Association for the Assessment and Accreditation of Laboratory Animal Care)- accredited facilities and were approved by the Institutional Animal Care and Use Committee (IACUC). Mice and hamsters were housed in the Mayo Clinic Animal Facility.

### Immunizations

Mice and hamsters were anesthetized with isoflurane prior to immunizations by the indicated routes: intramuscular (IM), intranasal (IN), sublingual (Sub). For intramuscular immunization, 50μL of a solution of virus diluted in PBS was injected into each flank for a total volume of 100μL per animal. Intranasal immunization was performed by pipetting 40μL of a solution of virus diluted in PBS dropwise into the nostrils of each animal. Each animal received a total volume of 40μL, alternating pipetting between nostrils.

### Bronchoalveolar Lavage (BAL)

BALs were performed as described in (*29*). Mice were euthanized via CO2 gas, then sterilized with 70% ethanol. Scissors were used to open the chest cavity up to the chin and to expose the trachea. A razor was used to puncture the trachea, and 1mL PBS was pipetted into and out of the lungs. This was repeated 2 additional times, giving a total volume of 3mL. Cells were then pelleted out of the bronchoalveolar lavage fluid by centrifugation. The cells were used to run flow cytometry, and the supernatant fluid was used to run ELISA.

### Lung Tissue Single Cell Suspension

Lung cells were isolated as described in (*29*). Briefly, lungs were extracted and processed with a gentleMACS dissociator (Miltenyi Biotec) and placed in gentleMACS C tubes containing 2.5mL RPMI, 40.4 μl of 14 U/mL concentration Roche Liberase TM, and 62.5 μl DNase I at 1 mg/mL concentration. Program Lung_01 was performed on the gentleMACS dissociator, followed by a one-hour incubation at 37°C. 2.5mL RPMI with 10% fetal bovine serum was added before running program Lung_02. All tubes were centrifuged for 5 minutes at 250 xg, and the contents were transferred to a 50 mL conical tube via a 70 μm mesh. 2.5 mL RPMI with 10% fetal bovine serum was added to the gentleMACS tubes and poured over the mesh to wash. 50 mL conical tubes were centrifuged for 10 minutes at 250 xg, and the supernatant was aspirated and discarded. Cell pellets were resuspended in 2 mL ammonium-chloride-potassium lysis buffer and centrifuged for 5 minutes at 350 xg. Supernatants from this reaction were discarded and the cell pellet was washed by resuspending in 2 mL PBS and was centrifuged for 5 minutes at 350 xg. Supernatants were extracted and discarded. The cell pellet was resuspended in desired volume RPMI and analyzed by flow cytometry.

### Sample Collections

At indicated time points, the animals were anesthetized with isoflurane and serum was collected by cheek bleed in mice or from jugular veins in hamsters. In addition, for bronchoalveolar lavage, the mice were euthanized via CO2 and bronchoalveolar lavage was performed according to the procedure described.

### Antibody ELISAs

Binding IgG and IgA antibody responses in mouse serum, hamster serum, and bronchoalveolar lavage fluid were measured by ELISA against spike S1 protein and SARS-CoV-2 receptor binding domain variants. Flat-bottom plates (ThermoFisher) were coated with 10 ng/well of spike S1 antigen in 100 μl PBS or 100 ng/well SARS-CoV-2 receptor binding domain variant antigen in 100 μl PBS, including a triplicate of negative control wells, which received no protein antigen. The protein antigen used for most SARS-CoV-2 ELISAs was recombinant SARS-CoV-2 (2019-nCoV) spike S1-Fc from Sino Biological. The protein antigen used for the SARS-CoV-2 variants included S1 proteins and receptor binding domain (RBD) Recombinant Proteins, also from Sino Biological. These included the following His6-tagged proteins expressed from 293 human cells: Wild-type; K417N; N429K; Y453F; S477N; E484K; N501Y; and K417N, E484K, N501Y, triple mutant corresponding to the South African variant, B.1.351. spike S1s with the single D614G and K417N, E484K, N501Y were also tested.

Plates were left overnight at 4°C. Plates were washed with 200 μl 1X PBS 2 times, followed by adding 200 μl per well of blocking buffer, consisting of 5% milk powder in TBST, for 2 hours at room temperature. Plates were washed with 200 μl 1X PBS 2 times. All samples were run in triplicate, including a triplicate of positive and negative control wells in each plate. Samples were serially diluted in blocking buffer and were transferred to the assay plate and incubated for 3 hours at room temperature. For the positive control wells, SARS-CoV-2 spike antibody [1A9] (GeneTex) was used as the primary antibody. Plates were washed with 200 μl 1X PBS 4 times, followed by addition of the secondary antibody. For hamster samples, the secondary antibodies used were: Peroxidase Conjugated Affinity Purified Anti-Golden Syrian Hamster IgG (H&L) Goat (Rockland Inc.), and Rabbit Anti-Hamster IgA (Brookwood Biomedical). For mouse samples, the secondary antibodies used were: GOXMO HRP HIGH XADS (Invitrogen), and HRP-Goat Anti-Mouse IgA (Invitrogen). The secondary antibody for the positive controls was Purified Recomb^TM^ Protein A/G Peroxidase Conjugated (Invitrogen). Plates were left to incubate with the primary antibody for 2 hours at room temperature. Plates were washed with 200 μl 1X PBS 4 times. 50 μl 1-Step^TM^ Ultra TMB-ELISA was added to each well and left at room temperature for 30 minutes, then 50 μl 2M sulfuric acid was added to each well. Plates were read at 450nm in a Synergy H1 microplate reader (BioTek). All statistical analyses were done by one-way ANOVA.

### Neutralization Assays

Pseudo-neutralization assays were performed on hamster serum using the cPass^TM^ Neutralization Antibody Detection kit (GenScript).

### ELISPOT Assay for Detecting Antigen-specific IFN-γ producing Cells

Freshly-isolated splenocytes were stimulated with spike S1 or S2 subunit protein (1 μg/mL) to determine the numbers of IFN-γ-producing cells by the Enzyme Linked Immuno Spot (ELISPOT) assay using the methodology reported previously (*30*). Briefly, splenocytes were plated at 2.5 × 10^5^ cells per well in triplicate in 96-well plates. Cells were incubated at 37 °C with medium alone, human papilloma virus E7 peptide (negative control), SARS-CoV-2 spike protein S1 subunit (1 μg/ml), SARS-CoV-2 spike protein S2 subunit (1 μg/ml), tetanus toxoid (TT, negative control, 100ng/ml)), or concanavalin A (Con A), positive control, 10ng/ml. After 24 h, cells were transferred to nitrocellulose plates, coated with anti-IFN-γ antibody, and incubated for 24 more hours. Plates were then washed and incubated with biotinylated anti-IFN-γ antibody, streptavidin-alkaline phosphatase, and colorimetric substrate, with washes between each step. After drying overnight, the plates were read on an AID ELIspot reader (San Diego, CA). Antigen-specific T cells were defined as the average number of spots elicited by the antigen of interest minus the average number of spots elicited when cells were incubated with culture medium alone, without the addition of any peptides.

### Flow Cytometry

Bronchoalveolar lavage samples were centrifuged at 1500rpm for 5 minutes, and the supernatant BAL fluid was removed. The cell pellet was resuspended in 200 μl T-Cell Media (IMDM with 10% FBS, Pen/Strep, and 2-ME), and was then added to a 96-well plate. 2 μl of stimulating mix (500 μg/mL ionomycin, 50 μg/mL PMA, and 445 μl T-Cell Media) was added to each well to be stimulated, and was incubated overnight at 37°C. 100 μl of a GolgiPlug Mix (1 μl GolgiPlug/1 mL T-Cell Media) was added to each well and thoroughly mixed by pipetting before incubating for 4-6 hours at 37°C. Cells were centrifuged at 1500rpm for 5 minutes and supernatant was removed. Cell surface antibodies were added in 50 μl total volume in FACS Buffer, then incubated 30 minutes on ice. Plates were washed twice with FACS buffer, resuspending cells in FACS Buffer and centrifuging for 5 minutes at 1500rpm to remove supernatant. The cells were then pelleted by centrifugation at 1500rpm for 5 minutes. Cell pellet was resuspended in 100 μl of Fix/Perm Solution for 20 minutes at 4°C before being washed twice with 1X BD Perm/Wash Buffer, then centrifuging at 1500rpm for 5 minutes to pellet. Intracellular cytokine antibodies were diluted in 50 μl 1X BD Perm/Wash Buffer and added to each well. Plate was then incubated 30 minutes on ice. Plates were washed twice with 1X BD Perm/Wash Buffer, resuspending cells in 1X BD Perm/Wash Buffer and centrifuging for 5 minutes at 1500rpm to remove supernatant. Cells were resuspended in 500 μl 1% PFA and left at 4°C overnight before flow cytometry analysis.

### Statistical Analysis

Prism 9 Graphical software was used for all statistical analyses.

## RESULTS

### Replication-defective and Single-cycle Adenoviruses Expressing SARS-CoV-2 Spike Protein

A codon-optimized cDNA encoding the original wild-type spike protein from the 2019-nCoV HKU-SZ-002a 2020 isolate was inserted into adenovirus vectors. This cDNA uses all native spike sequences and secretory leader and does not bear modifications such as proline mutations to alter spike structure (*4, 5, 31*). This cDNA was inserted into a cytomegalovirus (CMV) expression cassette and was used to generate human adenovirus serotype 6 (HAdV-C6, Ad6) vectors RD-Ad6-spike and SC-Ad6-spike, respectively (**Fig. 1A**). These vectors were tested for spike protein expression by infection of human A549 lung cells at varied multiplicities of infection (MOI) (**Fig. 1B**). Western blot on cells harvested 24 hours after infection demonstrated that both vectors produced spike protein; however, RD-Ad-Spike only generated detectable protein with 10^4^ virus particles (vp) per cell, but not with 100 vp/cell. In contrast, SC-Ad-Spike produced protein with 100 or more vp/cell with higher expression than RD-Ad at each dose.

**Fig. 1.**
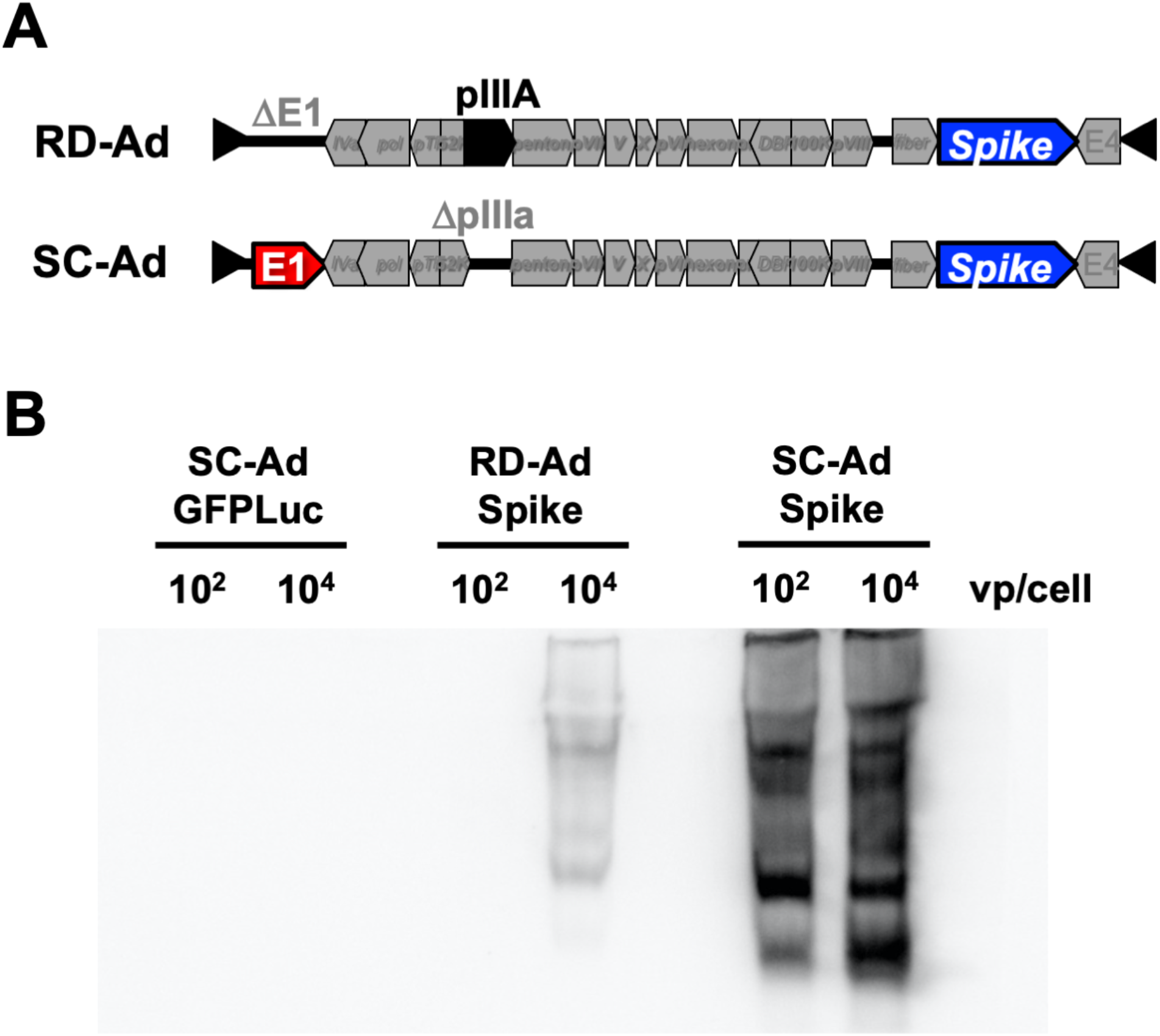
Replication-Defective and Single-Cycle Adenoviruses Expressing SARS-CoV-2 Spike. **A)** Schematics of replication-defective adenovirus (RD-Ad) spike vaccine construct as compared to single-cycle adenovirus (SC-Ad) spike vaccine construct. In the RD-Ad spike vector, the E1 protein has been removed and the SARS-CoV-2 spike protein has been inserted. In the SC-Ad spike vector, the pIIIA protein has been removed instead of the E1 protein, and the SARS-CoV-2 spike protein has been inserted. **B)** Western blot of human A549 lung cells infected with SC-Ad vector with GFP-Luciferase (SC-Ad GFPLuc), RD-Ad spike, or SC-Ad spike at doses of 10^2^ and 10^4^ viral particles per cell. Cells were harvested 24 hours post-infection, and western blot was performed looking for relative levels of SARS-CoV-2 spike protein.

### Comparison of Antibody Responses by RD-Ad-Spike and SC-Ad-Spike in Adenovirus-permissive Syrian Hamsters

Human Ad6 replicates its genome up to 100,000-fold in human cells (*32, 33*). SC-Ad replicates DNA identically to RC-Ad (*33*). Unfortunately, mice do not support the full life cycle or replication of human adenoviruses (*34*). Therefore, the administration of SC-Ad to mice may underrepresent the effect of transgene amplification that would be observed in humans and model organisms that are permissive to human adenovirus infection. In contrast to mice, Syrian hamsters are partially permissive for human adenoviruses (*34*) and their cells allow 350-fold replication of Ad6 DNA after infection (*24*). Immunization by RD-Ad and SC-Ad-Spike were therefore compared in Ad6-permissive Syrian hamsters to allow at least partial DNA replication to occur.

10^9^ vp of RD-Ad-Spike and SC-Ad-Spike were used to immunize male Syrian hamsters by IN and IM routes. These were compared to negative control RD- and SC-Ad expressing GFP-Luciferase (GL). Animals were immunized a single time and sera were collected at varied times thereafter (**Fig. 2**). Under these conditions, SC-Ad-Spike generated significantly higher spike antibody levels than RD-Ad-GL or SC-Ad-GL expressing GFP-Luciferase or when compared to RD-Ad-Spike at a fixed dilution (1/1,000) of sera. SC-Ad-Spike delivered by the intramuscular route generated higher IgG at all time points through 6 months after single immunization (p < 0.0001 by ANOVA) (**Figs. 2 and 3C and D**). SC-Ad-Spike by the intranasal route generated significantly higher IgG in sera at 6 weeks and 14 weeks, but not at 2 weeks and 6 months after single immunization (p < 0.0001 and p < 0.001 at 6 and 14 weeks). Serial dilution of these sera samples revealed reciprocal endpoint spike binding titers of 100 for RD-Ad-Spike by both routes (**Fig. 3A**). In contrast, SC-Ad-Spike by IN route had reciprocal binding titers of 1,000 (p < 0.05). SC-Ad-Spike by the IM route had reciprocal titers of 1,000 or greater (p < 0.0001). This experiment was repeated in female Syrian hamsters with similar results (**Supplemental Figs. 2 and 3**).

**Fig. 2.**
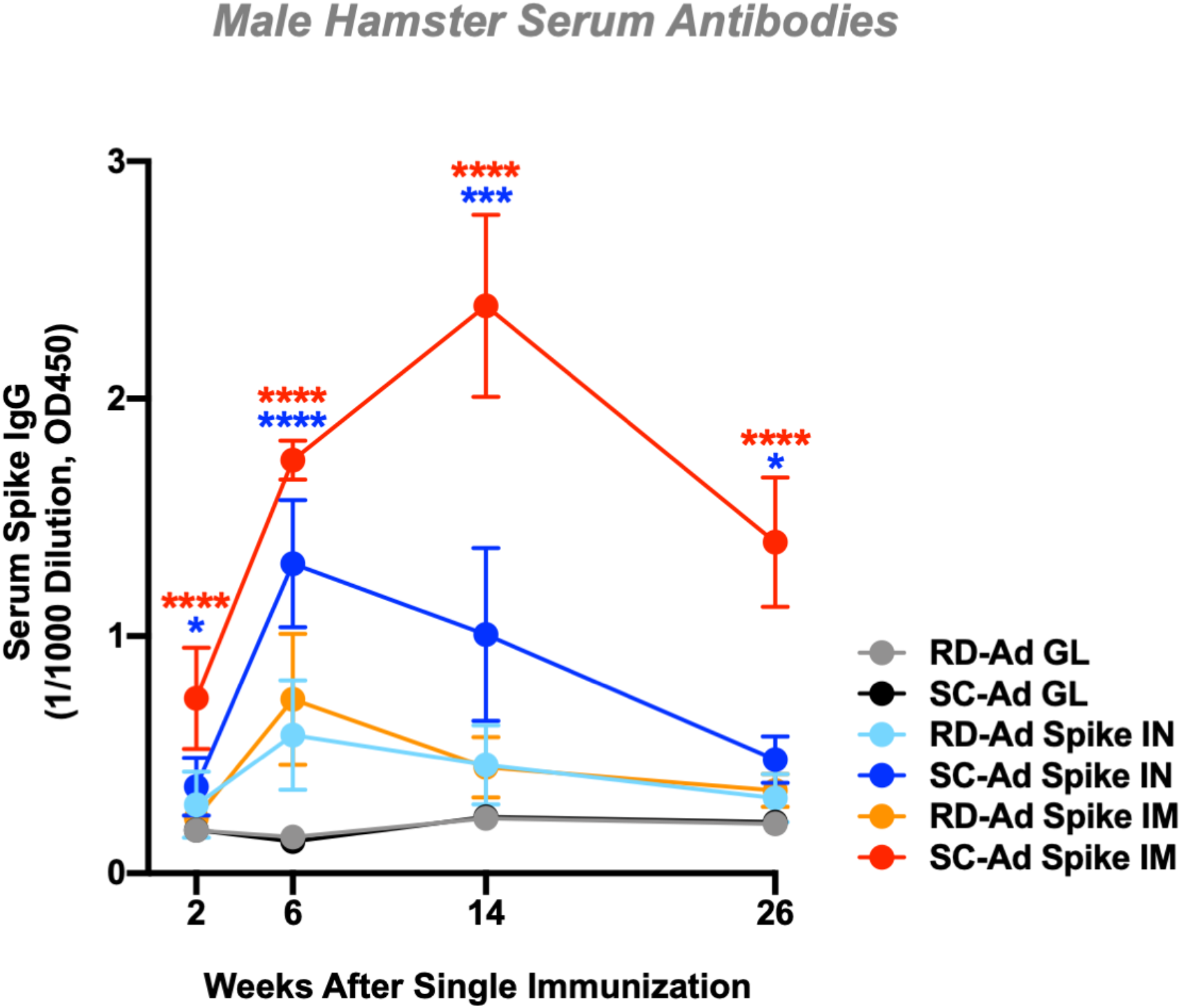
Kinetics of Spike Antibody Production by RD-Ad and SC-Ad Vaccines after a Single Intranasal or Intramuscular Vaccination. Male Syrian hamsters were immunized at a dose of 10^9^ vp, and serum was collected at weeks 2, 6, 14, 26 after single immunization. Serum was used at 1:1000 dilution to test for SARS-CoV-2 spike IgG antibodies by ELISA. (**** = p < 0.0001, *** = p < 0.0001, ** = p < 0.01, * = p < 0.05)

**Fig. 3.**
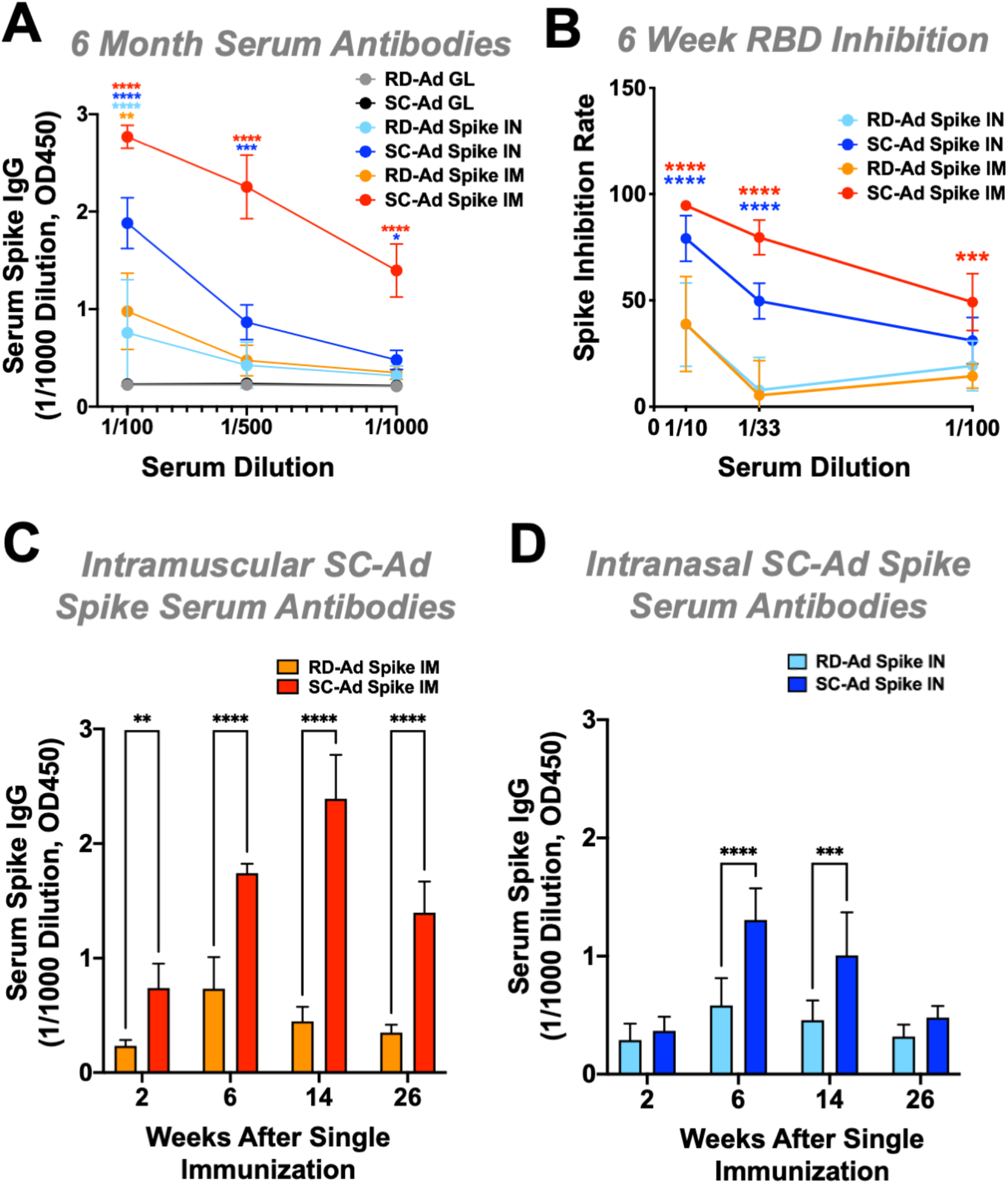
Spike Antibody Production by RD-Ad and SC-Ad Vaccines after a Single Intranasal or Intramuscular Vaccination. **A)** Male Syrian hamsters were immunized at a dose of 10^9^ vp, and serum was collected at 26 weeks (6 months) after single immunization. Serum was used at 1:100, 1:500, and 1:1000 dilutions to test for SARS-CoV-2 spike IgG antibodies by ELISA. **B)** Male Syrian hamsters were immunized and serum was collected at 6 weeks after single immunization. SARS-CoV-2 neutralization assay (Genscript) was performed at 1:10, 1:33, and 1:100 serum dilutions comparing RD-Ad spike and SC-Ad spike at intranasal and intramuscular routes of immunization. spike inhibition rate was determined based on the formula provided by Genscript. **C, D)** Comparison of serum spike IgG antibodies in Syrian hamsters immunized with RD-Ad spike and SC-Ad spike by intramuscular **(C)** and intranasal **(D)** routes of administration, analyzed by ELISA at 1:1000 serum dilution. (**** = p<0.0001, *** = p<0.0001, ** = p<0.01, * = p<0.05)

Varied dilutions of 6-week sera were assayed for SARS-CoV-2 neutralization antibodies with the cPass^TM^ Neutralization Antibody Detection kit that assays antibodies that block binding of spike receptor binding domain (RBD) to ACE2. Under these conditions, RD-Ad vaccinated hamsters failed to generate significant spike RBD inhibition at any dilution. In contrast, animals immunized with SC-Ad-Spike by the IN and IM route had significantly higher inhibition at all dilutions than RD-Ads within 6 weeks of single immunization (**Fig. 3B**).

### Antibody Binding to Spike Variants

SARS-CoV-2 has undergone rampant viral evolution as it has infected millions of humans. The emergence of SARS-CoV-2 variant B.1.1.7 in the UK, B.1.351 in South Africa, P.1 in Brazil, and a rash of other new variants raise significant concerns about the ability of vaccines to protect against them (*35–37*). These mutations are particularly concerning when they affect antibodies that bind to the ACE2 receptor binding domain, RBD, of spike. The SARS-CoV-2 variant B.1.1.7 contains H69del, V70del, Y144del, N501Y, A570D, D614G, and P681H mutations. The B.1.351 has K417N, E484K, N501Y, and D614G mutations. The P.1 variant bears K417T, E484K, and N501Y mutations (*35–37*). Mutations in the RBD domain are of most concern considering that they can impact the ability of neutralizing antibodies to block binding of the spike protein to ACE2.

Given concerns about these variants, week 14 sera from hamsters immunized IM with the negative control vaccine SC-Ad-GL and SC-Ad-Spike (**Fig. 2 and 3**) were compared for their ability to bind spike RBD (amino acids 319 to 541) and S1 variants (amino acids 16 to 685)(**Fig. 4 and Supplemental Fig. 3**). The RBD from the original SARS virus was also included as a reference. ELISAs performed at 1/1,000 dilutions demonstrated significant binding by samples from SC-Ad-Spike immunized animals to all the variant RBDs and S1 proteins when compared to SC-Ad-GL samples (p < 0.0001 by one-way ANOVA). SC-Ad-Spike bound the RBD from the original SARS-1 virus, but this did not reach significance at 1/1,000. When the samples were tested at 1/10,000 dilutions binding remained significant with p values remaining less than 0.0001 for all samples except the RBD with combined K417N, E484K, N501Y, which fell to a p value of less than 0.01. ELISA binding to the single E484K RBD was higher at this dilution than to other RBDs, suggesting some difference in structure or an artifact. When further 1/20,000 dilutions were tested, binding to all variants was still significantly different between SC-Ad-Spike and SC-Ad-GL samples (p < 0.05 to 0.0001), except for K417N, E484K, N501Y RBD, which was no longer significantly different. While binding to the K417N, E484K, N501Y RBD was lost, binding to the larger S1 protein with K417N, E484K, N501Y, and D614G mutations remained significant (p < 0.01). This was not unexpected, since the larger spike S1 protein has many more epitopes for polyclonal antibody binding.

**Fig. 4.**
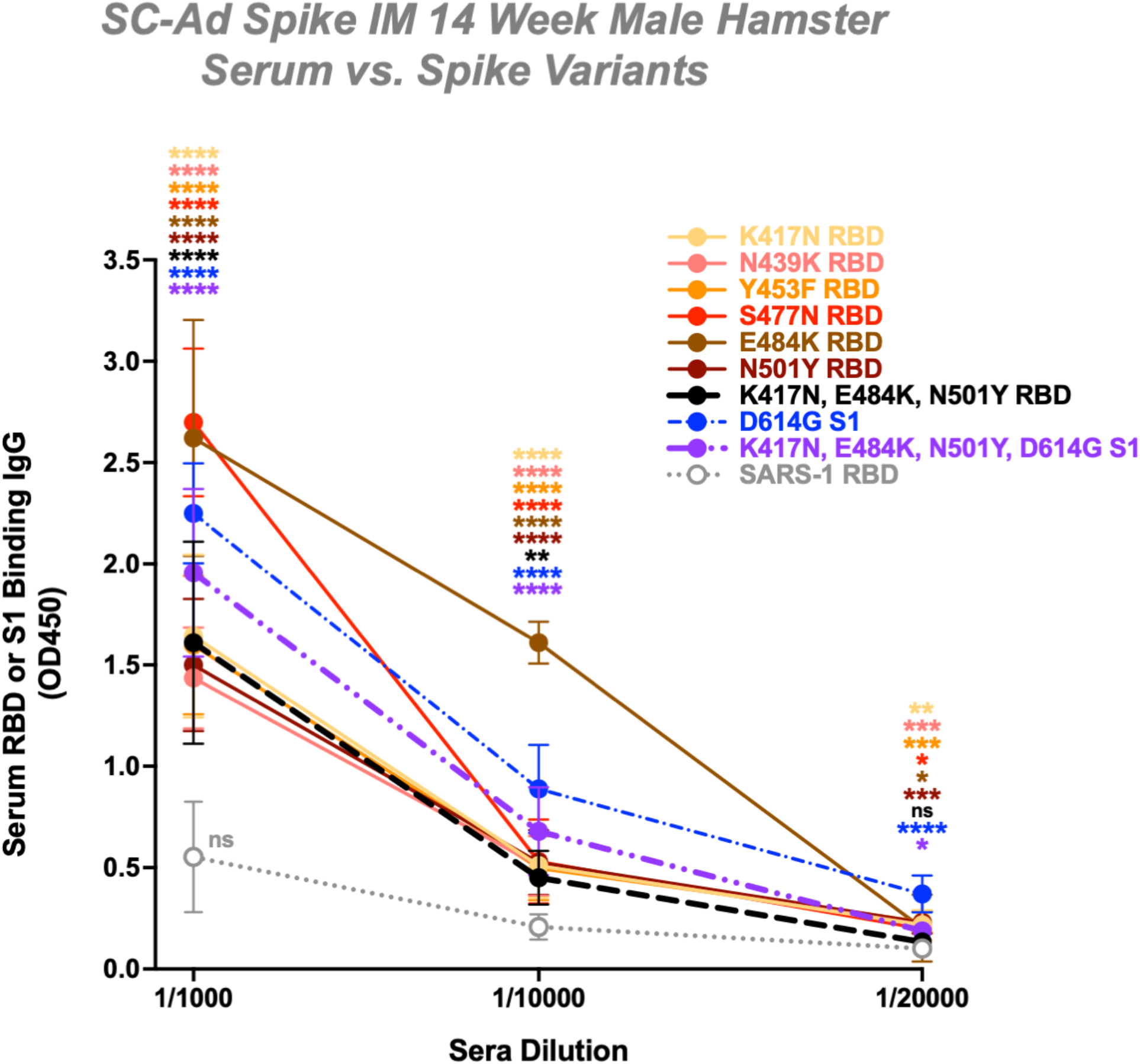
Antibody Binding to Spike RBD and S1 Variants. Week 14 sera from SC-Ad spike or SC-Ad-GL IM hamsters were analyzed for binding to variant RBDs and S1 proteins by ELISA at 1:1000, 1:10,000, and 1:20,000 dilutions. All levels of significance are shown as compared to the sample’s respective SC-Ad GL version. (**** = p<0.0001, *** = p<0.0001, ** = p<0.01, * = p<0.05). More detailed statistical comparisons are shown in Supplemental Fig. 3.

### Comparison of the Routes of Immunization by SC-Ad-Spike in Mice

Mice were utilized to evaluate the effects of the route of immunization of SC-Ad-Spike, since few immunological reagents exist to evaluate these responses in hamsters. BALB/c mice were immunized by IN and IM routes with PBS, 10^10^ vp of SC-Ad expressing Zika E protein, or 10^10^ vp of SC-Ad-Spike, and antibody responses were evaluated. This 10-fold higher dose was used to compensate for the lack of SC-Ad-Spike DNA replication in the mouse model. Under these conditions, mice immunized with SC-Ad-Spike generated robust IgG antibody responses within 2 weeks of immunization (p < 0.0001 by one-way ANOVA by both routes, **Fig. 5**).

**Fig. 5.**
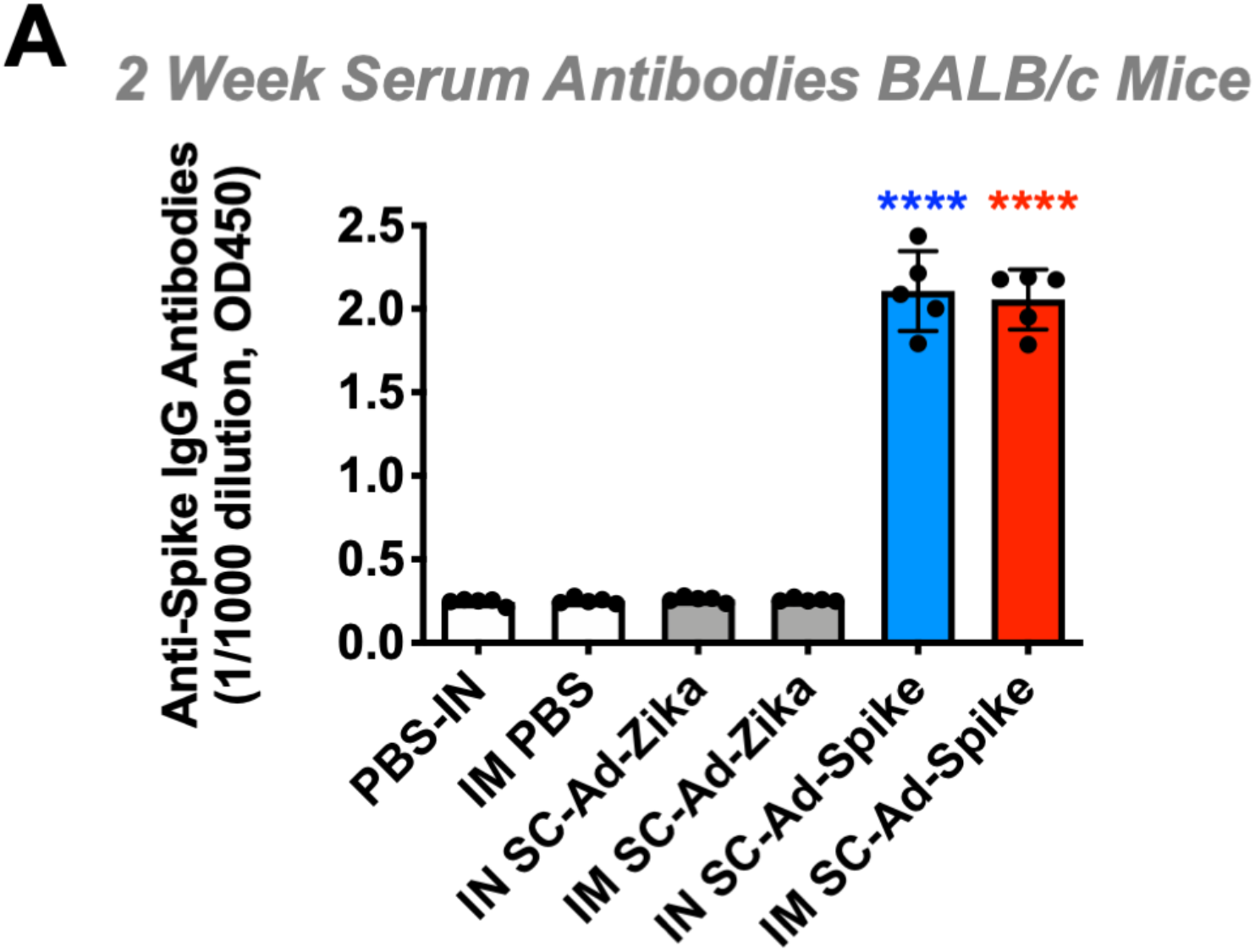
Serum Antibody Production to Spike Protein after Single Intranasal or Intramuscular Administration of SC-Ad-Spike in Mice. Male BALB/c mice were immunized at a dose of 10^10^ vp virus, and serum was collected 2 weeks after single immunization and was tested for SARS-CoV-2 spike IgG antibodies by ELISA. (**** = p<0.0001, *** = p<0.0001, ** = p<0.01, * = p<0.05)

### Effects of the Routes of Immunization on Mucosal Antibody Responses in the Lungs

Five animals from selected groups in **Fig. 5** were sacrificed 8 weeks after single immunization and bronchoalveolar lavages (BALs) were performed to collect mucosal antibodies and immune cells from the lungs. The 3 mL BAL washes were diluted 1/500 and were used to detect anti-spike IgG and IgA antibodies by ELISA. By 8 weeks after single immunization, the mice had significant levels of IgG antibodies in their BALs (p < 0.01 and 0.001, **Fig. 6A**). Notably, significant anti-spike IgA antibodies were only observed in IN-immunized mice (p < 0.01). When BAL samples were tested for RBD neutralizing activity, both the IN and IM-immunized mice had significant activities (p < 0.01 and 0.05, respectively, **Fig. 6B**).

**Fig. 6.**
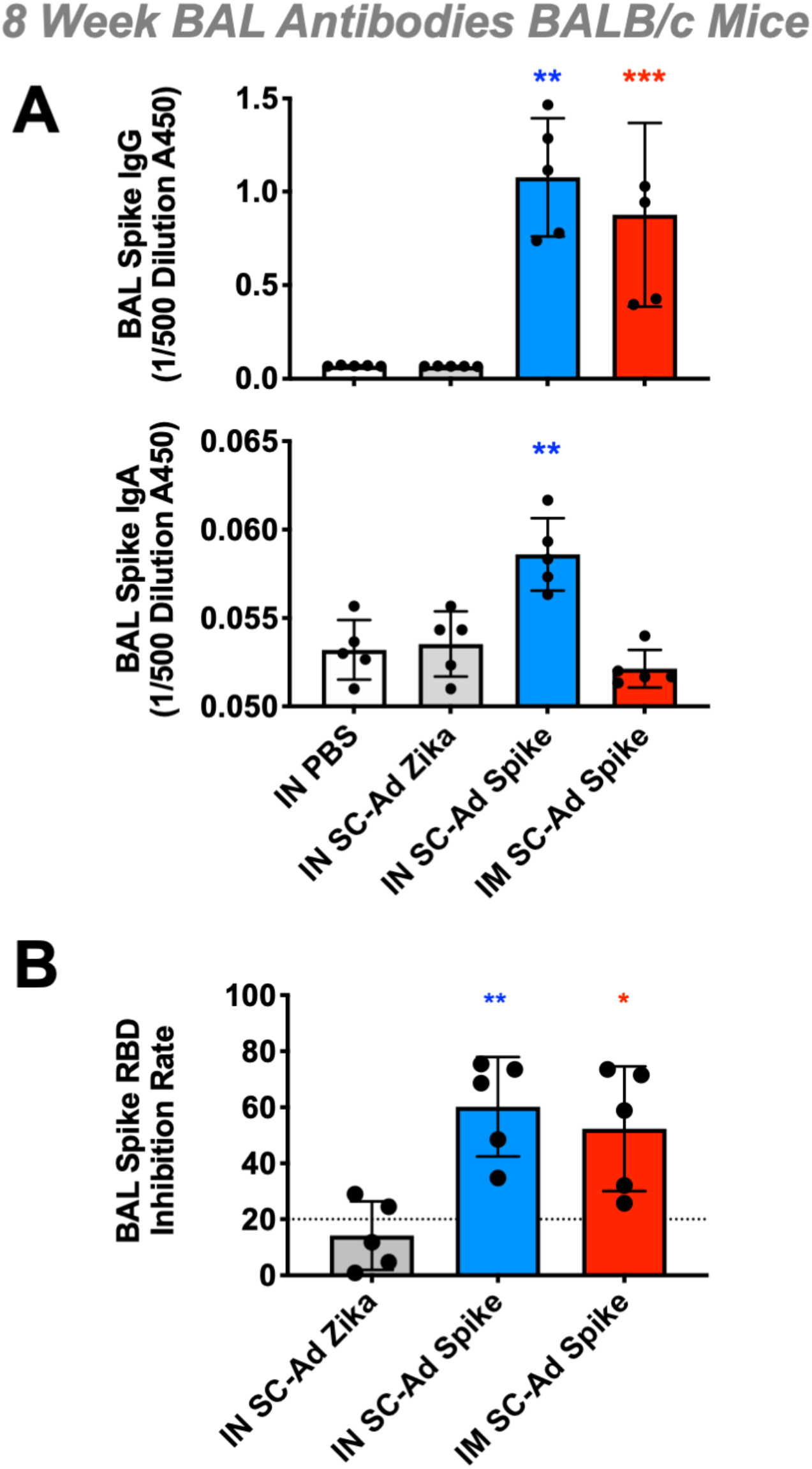
Mucosal Antibody Production to Spike Protein in Bronchoalveolar Lavages (BALs) after Single Intranasal or Intramuscular Administration of SC-Ad-Spike in Mice. Male BALB/c mice were immunized with 10^10^ vp of SC-Ad-Spike and bronchoalveolar lavage (BAL) fluid was collected at 8 weeks after single immunization. **A)** BAL fluid was used at 1:500 dilution to test for SARS-CoV-2 spike IgG and IgA antibodies by ELISA. Plates were read at 450nm, and all analyses were done by one-way ANOVA. **B)** SARS-CoV-2 neutralization assay (Genscript) was performed at 1:10 dilution of BAL fluid, comparing IN SC-Ad Zika, IN SC-Ad spike and IM SC-Ad spike. spike inhibition rate was determined based on the formula provided by Genscript. (**** = p<0.0001, *** = p<0.0001, ** = p<0.01, * = p<0.05).

### Effects of the Routes of Immunization on Mucosal T Cell Responses in the Lungs

BAL samples were also examined for the presence of T cells in the lung. The small number of cells obtained and an absence of known spike T cell epitopes for BALB/c mice precluded testing for spike-specific responses; however, when flow cytometry was performed, IFN-γ and IL-4-expressing CD4 and CD8 T cells were detected in BAL samples (**Fig. 7**). These analyses revealed no significant increases in CD4 or CD8 T cells in BAL samples after IM immunization. In contrast, there were significant increases in CD8^+^ IFN-γ^+^, CD4^+^ IFN-γ^+^, and CD4^+^ IL-4^+^ T cells in the BALs of animals immunized intranasally with either SC-Ad-Zika E or SC-Ad-Spike.

**Fig. 7.**
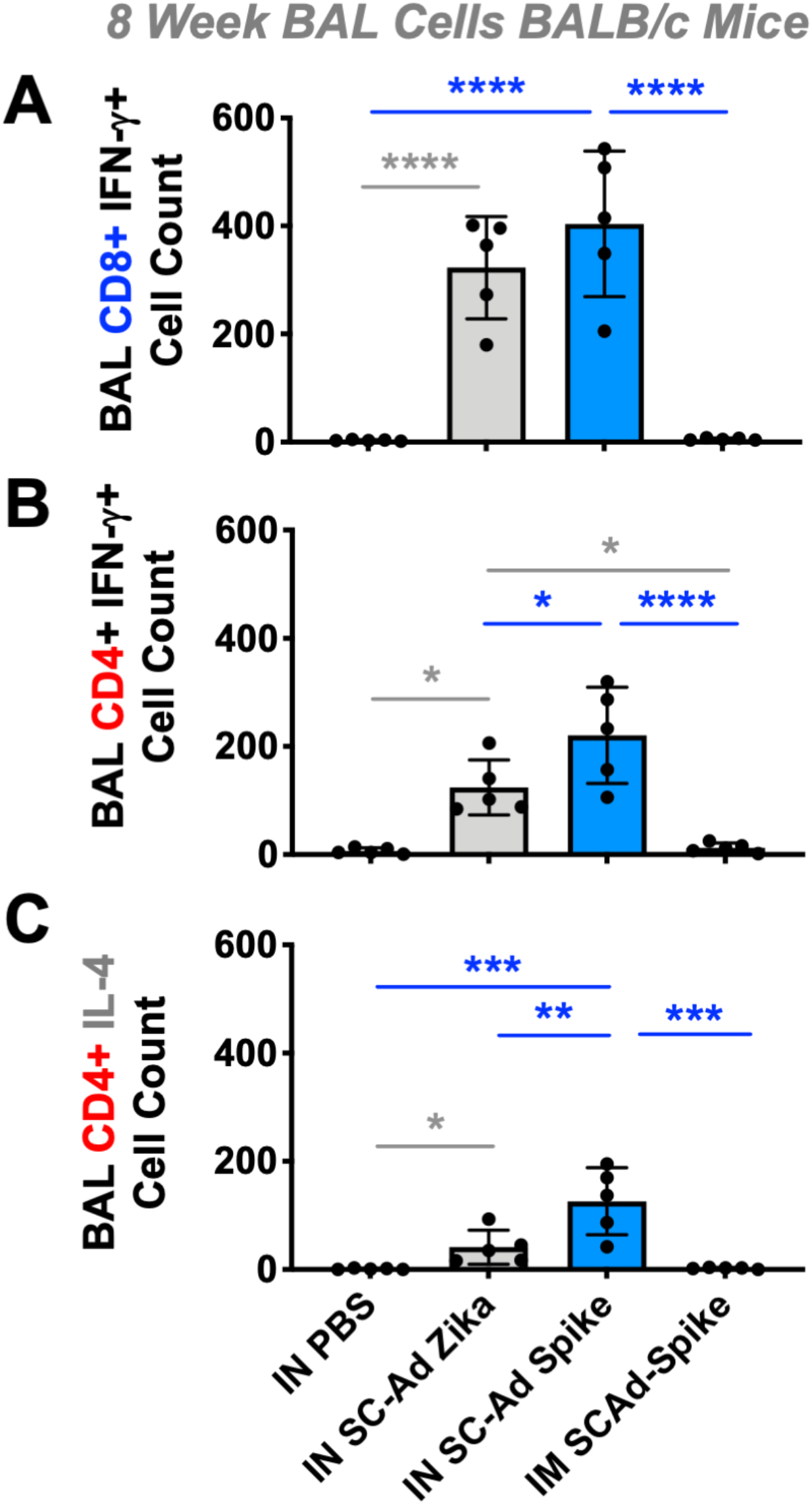
Changes in T Cell Populations in Bronchoalveolar Lavages (BALs) after Single Intranasal or Intramuscular Administration of SC-Ad-Spike in Mice. 8 weeks after single immunization of 10^10^ vp of SC-Ad-Spike in BALB/c mice, BAL was performed. Cells were pelleted out of BAL fluid and analyzed by flow cytometry. **A)** Shows number of CD8 T cells counted that expressed IFN*γ*. **B)** Shows number of CD4 T cells counted that expressed IFN*γ*. **C)** Shows number of CD4 T cells counted that expressed IL-4. (**** = p<0.0001, *** = p<0.0001, ** = p<0.01, * = p<0.05).

### Effects of the Routes of Immunization on Systemic T Cell Responses

Splenocytes from the animals in **Fig. 7** were assayed for IFN-γ, IL-17, and IL-4-expression by ELISPOT after stimulation with spike S1 whole protein subunit (**Supplemental Fig. 4**). These data demonstrated higher splenocyte IFN-γ, IL-17, and IL-4 responses in the BALB/c mice after IN immunization than after IM immunization.

## DISCUSSION

The purpose of this study was to compare RD- and SC-Ad vaccines expressing the SARS-CoV-2 spike protein and to evaluate whether mucosal immunization may have utility when considering vaccines against a mucosal pathogen like SARS-CoV-2. These data suggest that vaccines that retain the ability to replicate their DNA can drive more potent and long-lasting immune responses than non-replicating vaccines. These data also suggest that there may be advantages to delivering vaccines at mucosal sites by single immunization or as a prime followed by later boost.

Consistent with previous comparisons, SC-Ad expressed higher levels of antigen than matched RD-Ad vector. This higher expression by SC-Ad-Spike translated into higher serum antibody responses than RD-Ad-Spike in adenovirus-permissive Syrian hamsters after single immunization by either the IM or IN route. The levels of antibodies in sera were higher in animals immunized by the intramuscular route than the intranasal route. In all the animal models, antibody titers were higher in ELISA assays than in varied neutralization assays, as expected. This may reflect, in part, the use of the wild-type spike protein sequence rather than modified spikes locked in the “up” position (*4*). Use of such a modified spike would likely increase neutralization by SC-Ad.

SC-Ad-Spike generated antibodies from these hamsters were able to cross-react in ELISA assays against several single point mutant RBD variants, including those observed in the U.K. B.1.1.7 strain and the South African B.1.351 strain. These antibodies were also able to bind K417N, E484K, N501Y RBD at 1/1,000 and 1/10,000 dilutions in ELISA, but were insignificant at 1/20,000 dilutions. This was not surprising given that the presence of three separate mutations have been shown to affect the ability of Pfizer, Moderna, and other vaccines to recognize these new variants.

The single-cycle replication “engine” can be applied to any adenovirus serotype. This increased potency could be utilized in two ways. In one, SC-Ad is delivered at the same virus particle doses as current RD-Ad COVID-19 vaccines to garner stronger immune responses. In the second, SC-Ad is used at a lower dose, perhaps 10 to 100-fold lower, to deliver equal potency to RD-Ad vaccines, but allowing 10 to 100 times more vaccine doses from the same size of a GMP vaccine production run. This could be pivotal for expanding access to vaccines for this pandemic or the next to vaccinate people in rich and poorer countries.

Species C human Ad6 was used to test proof of concept since it is equal to or is more robust than species C Ad5 as a vaccine or as an oncolytic (*32, 38–42*). Species C Ad5 and Ad6 also appear to be more robust as gene-based vaccines than lower seroprevalent viruses like species D Ad26 and species E ChAdOx1 (*38, 39, 43, 44*).

Another interesting aspect of this work was examining if there is utility in applying these gene-based vaccines at mucosal surfaces. Hamsters have few immune reagents, so this was examined in more detail in mouse models. In mice, we show that intranasal immunization generated equal IgG antibodies in the lungs of mice, but higher IgA and CD4 and CD8 T cells in BAL samples. Intramuscular immunization was able to generate IgG antibodies in BAL fluid. However, the IM route reduce markedly low IgA antibodies than the IN route. Likewise, IM immunization failed to traffic CD4 or CD8 T cells to the lumen of the lung in contrast to IN mucosal immunization. These data suggest that IN mucosal routes of immunization may do a better job at placing effector antibody and T cells at the sites of earliest exposure to SARS-CoV-2.

These observations are consistent with our previous observations testing IN and IM prime-boosts with SC-Ad expressing HIV envelope in rhesus macaques (*26*). Animals immunized with SC-Ad by only the IM route had lower HIV-1 antibody-dependent cellular cytotoxicity (ADCC) antibody activity and lower levels of peripheral T follicular helper (pTfh) cells in their lymph nodes. Conversely, animals immunized with SC-Ad by the IN route had higher ADCC, higher Tfh cell counts in lymph nodes, and lower SHIV viral loads in their gastrointestinal tracts after rectal SHIV challenge (*26*). These data suggest that there may be benefits in priming the immune system at mucosal sites when immunizing against pathogens that also enter at these sites.

Mucosal immunization may also have utility to impact vaccine safety. There are concerns with observations of thrombotic thrombocytopenia in a small number of people who have received the Ad26 and ChAdOx1 COVID-19 (*45, 46*). These vaccines were delivered by the intramuscular route. Adenoviruses do not naturally infect the muscle. Ads naturally infect humans at some mucosal site (reviewed in (*47*)). Injection of anything into the muscle can breach blood vessels and allow leak into the bloodstream. For an intramuscular Ad vaccine, this can cause adenovirus to be absorbed by liver Kupffer cells, liver sinusoidal endothelial cells, and to productively infect liver hepatocytes, spleen, or lungs. It is therefore possible that injecting Ads into the muscle by this unnatural route may elevate the risk of side effects. It is possible that delivering adenovirus vaccines by intranasal vaccination may avoid some of these new risks. That having been said, it should be noted that intranasal immunization has its own possible side effects including retrograde transport into the olfactory bulb and Bell’s palsy.

Another consideration is the use of adenoviruses that are low seroprevalent in humans. The primary advantage of Ad26 and ChAdOx1 is that few people have been exposed them naturally and so most people do not have neutralizing antibodies against these vaccines. While that is true, it also means that there is less experience with how these viruses behave in humans and what side effects may be observed.

In contrast, 27 to 100% of humans are immune to Ad5 (*43*). Ad6 is lower seroprevalent than Ad5, with 4 to 22% of humans having already been exposed to Ad6 (*48, 49*). In both cases, these humans have experienced these species C Ads after mucosal exposure without obvious connections to side effects associated with COVID-19 vaccines. One might say that both Ad5 and Ad6 have been field tested as mucosal vaccines in as many as a billion humans. Conversely, no humans have been naturally exposed to Ad5, Ad6, Ad26, or ChAdOx1 by intramuscular exposure.

While these common human adenoviruses may have some ironic safety value, it is still possible that arming any adenovirus or any gene-based vaccine with the SARS-CoV-2 spike protein may by itself play a role in the observed thrombotic side effects by intramuscular or mucosal routes of vaccination.

These concepts need to be explored further. Regardless, this study suggests that mucosal immunization may have value when combating SARS-CoV-2 and other mucosal pathogens. This work also suggests that giving adenovirus vaccines the ability to replicate via single-cycle modifications may have value in increasing per virus potency or by allowing more doses to be produced by using fewer virions per person.

## Supporting information

Arrive 10 checklist

## Acknowledgements

This work was funded by grants from Fastgrants.org/Mercatus and from the Harrington Foundation COVID-19 Scholar Program to M.A.B. This project was supported by the Walter & Lucille Rubin Fund in Infectious Diseases Honoring Michael Camilleri, M.D. at Mayo Clinic. This project would not have been possible without generous support from Mayo Benefactors and the Mayo Clinic COVID-19 Task Force.

