## Supplemental Figures for "Comparison of Mucosal and Intramuscular Immunization against SARS-CoV-2 with Replication-Defective and Replicating Single-cycle Adenovirus Vaccines"

**Supplemental Figure 1    Mudrick *et. al.***

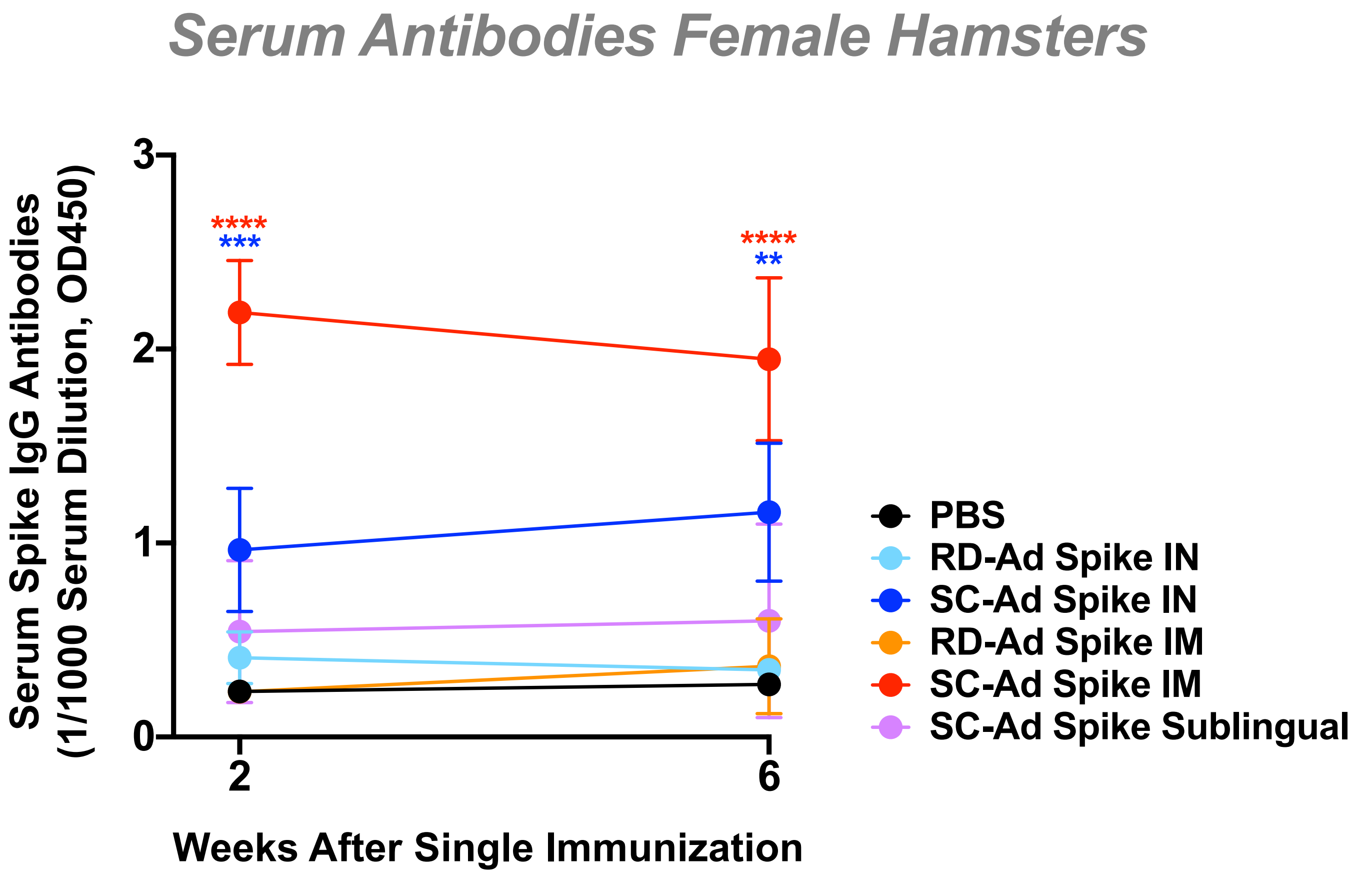


**Supplemental Figure 1. Spike Antibody Production by RD-Ad and SC-Ad Vaccines after a Single Intranasal or Intramuscular Vaccination in Female Hamsters.** Female Syrian hamsters were immunized at a dose of $10^9$ vp, and serum was collected at indicated times and tested t  1:1000 dilutions to test for SARS-CoV-2 spike IgG antibodies by ELISA. (**** = p<0.0001, *** = p<0.0001, ** = p<0.01, * = p<0.05)

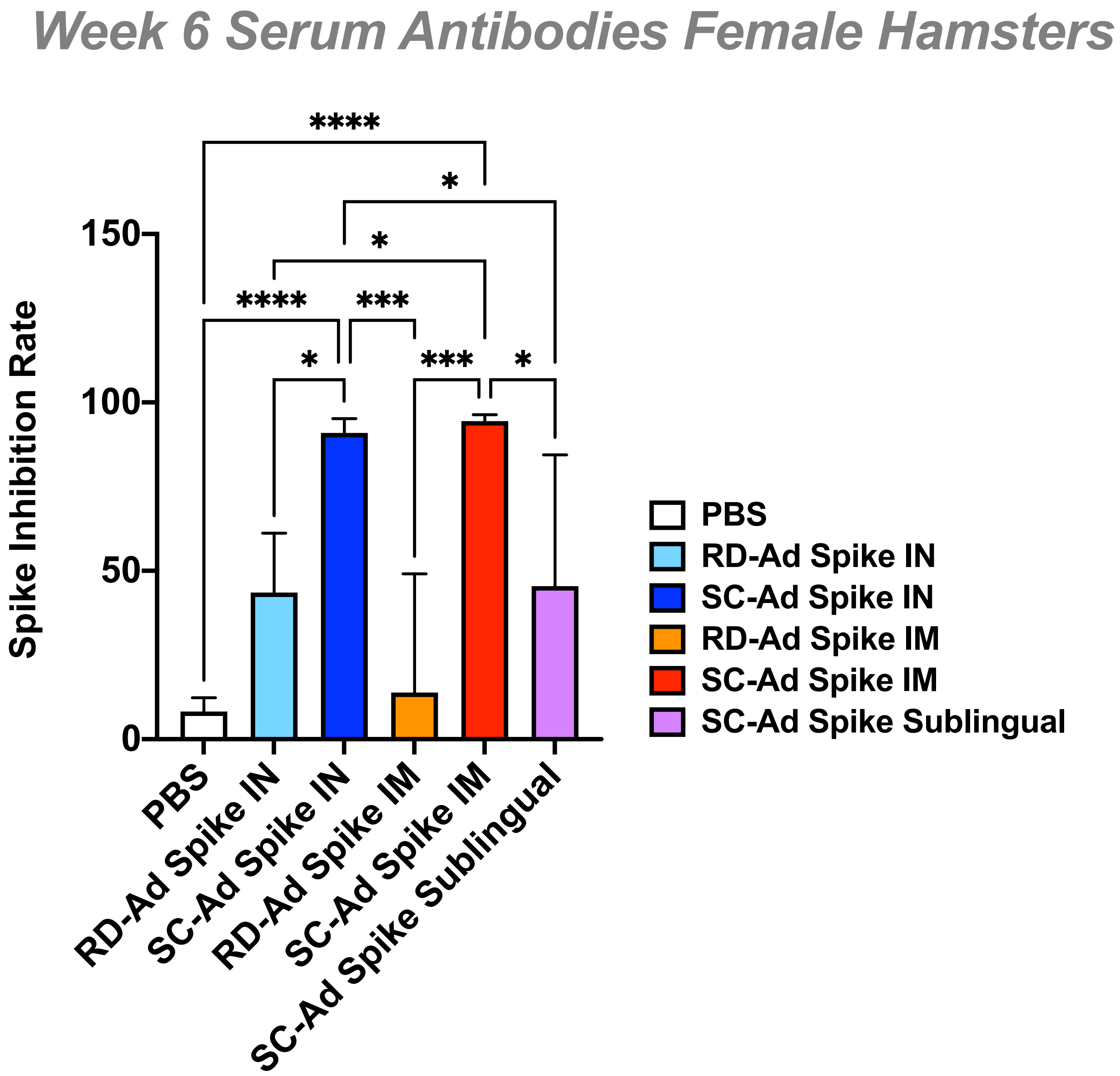


**Supplemental Figure 2. Spike Antibody Production by RD-Ad and SC-Ad Vaccines after a Single Intranasal or Intramuscular Vaccination in Female Hamsters.** Female Syrian hamsters were immunized at a dose of $10^9$ vp, and 6 week serum ELISAs were compared by ANOVA. (**** = p<0.0001, *** = p<0.0001, ** = p<0.01, * = p<0.05)

### Supplemental Figure 3 Mudrick *et. al.*

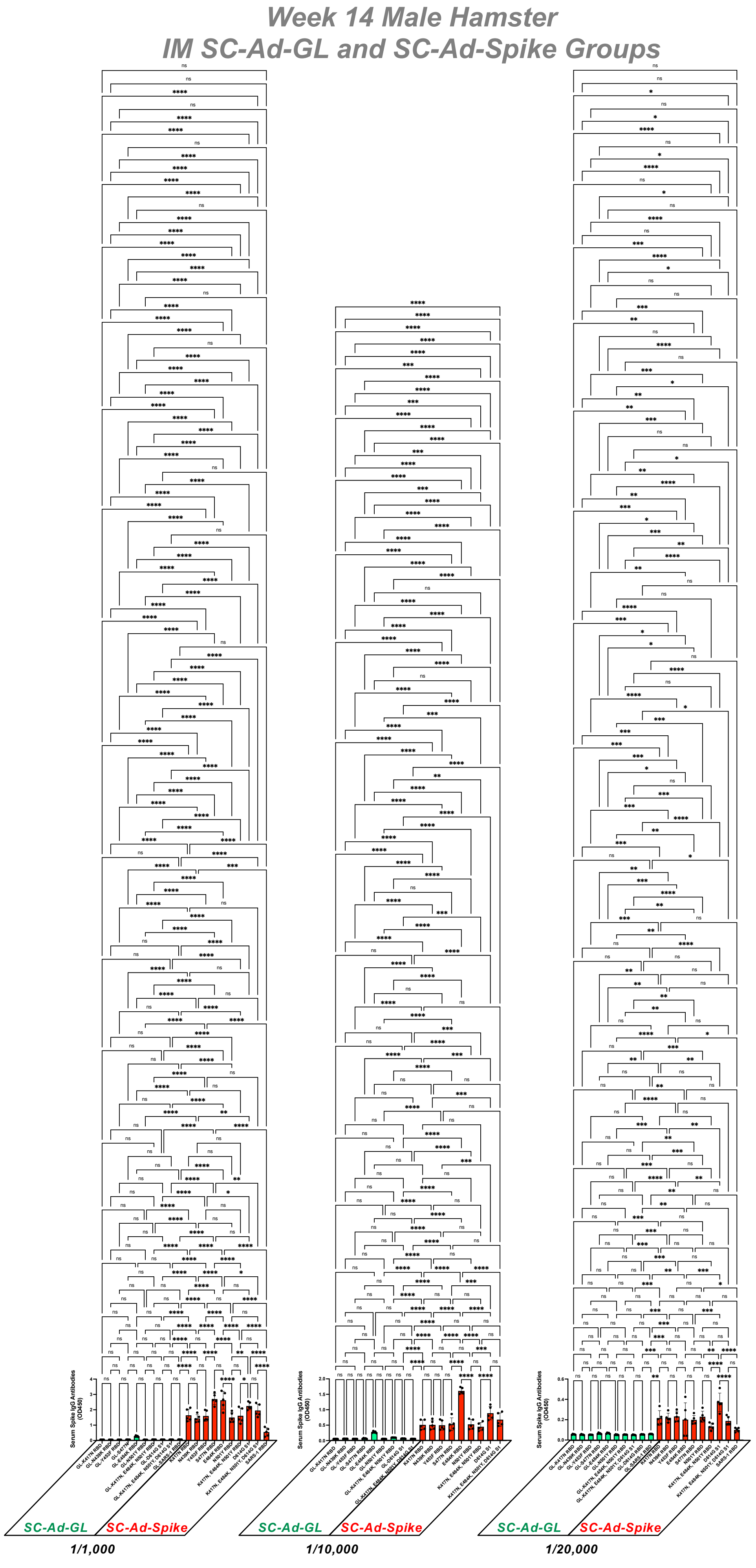


**Supplemental Figure 3. Full ANOVA Comparisons of RD-Ad and SC-Ad Vaccines after a Single Intranasal or Intramuscular Vaccination in Male Hamsters.** Data from Figure 2. (**** = p<0.0001, *** = p<0.0001, ** = p<0.01, * = p<0.05).

Supplemental Figure 4    Mudrick *et. al.*

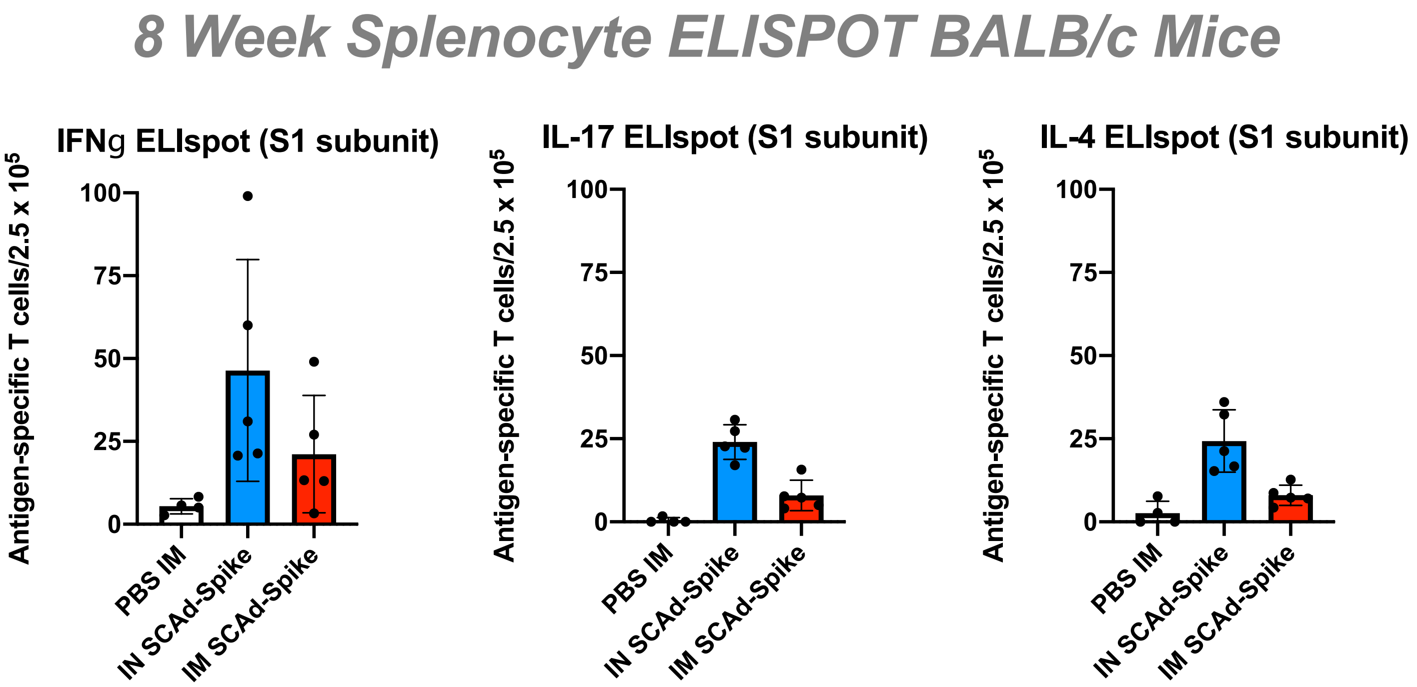


**Supplemental Figure 4. ELIspot Responses in Splenocytes after a Single Intranasal or Intramuscular Vaccination in Mice.** BALB/c mice were immunized by the indicated routes with $10^{10}$ vp of SC-Ad-Spike or PBS. Splenocytes were stimulated with whole S1 spike subunit and IFNγ ELIspot was performed (* $p<0.01$).
